# Unravelling the ageing-reversal potency of stem cell-derived extracellular vesicles in a rat model of premature cardiac senescence

**DOI:** 10.1101/2024.01.28.577660

**Authors:** L Gómez-Cid, A Campo-Fonseca, M Cervera-Negueruela, A Ocampo, A Pinto, JM Gil-Jaurena, S Suárez-Sancho, F Fernández-Avilés, J Bermejo, L Grigorian-Shamagian

## Abstract

Cardiosphere-derived cells (CDCs) and the extracellular vesicles they release (CDC-EVs) have demonstrated to induce rejuvenation and to improve cardiac structure and function, but their efficacy may vary among donors and there is still a lack of adequate potency assays. We aimed to identify parameters that could easily predict the ageing-reversal potency of CDC-EVs. CDCs derived from cardiac tissue of 34 human donors (age range: 3-months to 81-years old) were characterized in terms of their phenotypical and biological properties. The anti-senescent activity of the CDC-EVs was assessed *in vitro* using a predesigned matrix assay with several ageing-related markers. We found that while CDC-donor’s chronological age was not determinant, the degree of CDC senescence in culture showed a strong correlation to most CDĆs bioactive properties. However, CDC senescence alone was insufficient to predict CDC-EV *in vitro* potency. Thus, we scored the potency of the CDC-EVs based on their performance in the newly designed *in vitro* matrix assay and classified them as more-(P-EVs) and less-potent (NP-EVs). We then tested P- and NP-EVs in a rat model of premature cardiac ageing and found that only the P-EVs reduced senescence-associated *GLB1* gene expression in the heart and tended to protect it from hypertrophy development and fibrosis. On the contrary, NP-EVs failed to induce any improvements and negatively affected cardiac hypertrophy, fibrosis, and perfusion. In conclusion, despite CDC-EV anti-ageing potency cannot be predicted by the chronological age of the donors or CDC senescence as surrogate markers, we propose an *in vitro* potency assay that could be used to evaluate allogenic EV suitability before its use in the treatment of cardiac ageing.

## Introduction

Cellular, structural and functional changes associated to ageing are largely responsible for heart diseases such as heart failure with preserved ejection fraction (HFpEF) (Steenman & Lande, 2017), an unmet medical need because of increasing prevalence, high mortality and morbidity rates, and limited therapeutic strategies (Borlaug, 2020; Redfield & Borlaug, 2023). Cardiac ageing is associated to increased cell senescence (Abdellatif et al., 2023; Climent et al., 2017), compromised myocardial perfusion (Uren et al., 1995), and increased fibrosis and hypertrophy (Gazoti Debessa et al., 2001) among others. Cardiosphere-derived cells (CDCs) and the extracellular vesicles they release (CDC-EVs) present promising characteristics for reverting some of these ageing-associated pathological features as demonstrated in preclinical studies (Grigorian Shamagian et al., 2023; Grigorian-Shamagian et al., 2017), such as increasing microvessel formation (Malliaras et al., 2014), attenuating cardiomyocyte (CM) apoptosis and hypertrophy, reducing regional fibrosis (Chimenti et al., 2010; Tseliou et al., 2014) and acting as immunomodulators (López et al., 2020; Tseliou et al., 2014).

However, when moving stem cells or extracellular vesicles as products from preclinical to clinical scenario the results are less satisfactory than expected (Grigorian-Shamagian et al., 2021). Hasty translation led to underestimate how the heterogeneity and complexity of these products could impact their potency (Povsic et al., 2021), one of the factors that could influence the disappointing results of previous clinical trials. Despite most clinical trials in the field used confirmation of cell identity as the single requirement before product administration (Gómez-Cid et al., 2021), the gold standard for cell potency assessment in cardiac applications is structural and functional recovery in a rodent model of myocardial infarction (Cheng et al., 2014; Ibrahim et al., 2019). However, this model is costly, time-consuming and of doubtful translation, hindering its use for routine testing (Porat et al., 2015). *In vitro* functional assays do not reflect the complexity of living organisms, but they require less time and economic resources and can be representative of the potential mechanism of action (MoA) of the tested product if properly validated. Finding economic, feasible and efficient potency assays based on evidence of MoA for therapeutic cells and cell-derived EVs in the cardiovascular field still remains a challenge (Gómez-Cid et al., 2021; Marbán, 2018; Willis et al., 2017), and crucial for successful translation.

Considering relevant characteristics and the potential MoA by which CDC-EVs may exert their beneficial effects in the treatment of cardiac ageing, we here explore if the donorś chronological age, senescence of the CDCs and/or CDC-EV rejuvenating potency *in vitro* can be used to predict CDC-EV efficacy *in vivo*.

## Methods

### Characterization of Cardiosphere-derived cells and extracellular vesicles

Study protocol was approved by the local ethics committee for human research and all patients, or their legal guardians signed the written consent before inclusion in the study. Heart tissue was collected from 34 patients who underwent cardiac surgery for other reasons. Patients of both sexes and different ages (including pediatric and adult) were randomly included in the study. Donorś age and sex distribution are detailed in Suppl. Table 1 and Suppl. Figure 1.

CDCs and CDC-EVs were subsequently obtained following the procedure published previously (Gómez-Cid et al., 2020). CDC identity was confirmed using cell surface markers and CDC-EV concentration and size quantified through Nanoparticle Tracking Analysis (NTA) (Suppl Figure 2). The CDCs were characterized using genetic expression profile, biological age and activity and CDC-EV were characterized in terms of their bio-activity using an ageing-reversal potency matrix assay.

### Design of the ageing-reversal potency matrix assay

The CDC-EV rejuvenating and pro-angiogenic potency was tested *in vitro* on human cardiac stromal cells (CSCs) and human endothelial cells (ECs). Two types of CSCs were used as target: moderately senescent (25.6 ± 1.4 % of senescent cells in culture) from donor 1 (54 years old, male) and highly senescent (47.5 ± 1.3 %) from donor 2 (56 years old, female). The rejuvenating potency of CDC-EVs was determined by their ability to reduce CSC senescence, to induce CSC IL-6 secretion and to reduce the genetic expression of *CDKN1A* (p21), *CDKN2A* (p16), *TP53* (p53) and *TGFB1* (TGF-β), senescence and fibrosis-related genes in CSCs. CDC-EVs pro-angiogenic potency was determined by their ability to induce tube formation on human umbilical vein EC.

The results of all the tests in the matrix-assay for each CDC-EVs were scored individually and added to estimate the global potency score as shown in Suppl. Table 2. The CDC-EVs with the highest total score were selected as more-potent (P-EV) and the CDC-EVs with the total lowest score were selected as less-potent (NP-EV). The maximum possible score was 18 if CDC-EVs showed superior performance in all the tests in the matrix-assay.

### In vivo experiments in a rat model of premature senescence

All methods were carried out in accordance with relevant guidelines, approved by the Institutional Animal Care and Use Committee of the Gregorio-Marañon University Hospital. The experimental procedure is illustrated in Figure 4a. Thirty-six 12-week-old Sprague-Dawley rats (30% females) were randomly allocated into four groups: healthy controls (*n* = 11), D-GAL controls (sham group, D-gal, *n* = 12), D-GAL rats treated with more potent CDC-EVs (D-gal + P-EVs, *n* = 7) and D-GAL rats treated with less-potent CDC-EVs (D-gal + NP-EVs, *n* = 6). To induce cardiac ageing, animals in the D-GAL groups received during 13 weeks daily intraperitoneal (IP) injections of 600 mg/kg of D-galactose (D-gal, G53688, Sigma Aldrich) resuspended in phosphate buffered saline (PBS) at 300 mg/ml as described (Bo-Htay et al., 2018). For 13 weeks, animals in the healthy control group received daily IP injections of PBS alone. After 13 weeks, rats in the D-GAL group were randomly allocated to receive IP administration of: 1) PBS alone; 2) P-EVs resuspended in PBS; or 3) NP-EVs resuspended in PBS. The dose of P- and NP-EVs was of 7.5 μg EV protein/g animal weight. During the following 7-weeks, the animals received additional IP injections of D-GAL or PBS alone twice a week based accordingly to their allocation group. In a randomly selected subgroup of rats, cardiac perfusion was evaluated using Single Photon Emission Computer Tomography (SPECT) at 12 weeks (n=6) and 19 weeks (n=18). At the endpoint, the animals were euthanized, blood samples were collected, and hearts were excised for further studies.

### Statistical analysis

Results are presented as mean ± standard error of the mean in the text and in figures. Continuous variables were compared under the different conditions using Student’s t-tests. Categorical variables (percentage of senescent cells and of Ki-67+ cells) were compared using the Z-Score for two population proportions. Pearson tests were performed to study the correlation among the different parameters. All probability values reported are two-sided, with *p* < 0.05 considered significant. For *in vitro* studies the lowest number of replicates per experiment was three.

## Results

### The senescence status more than the chronological age of cell donors determines the cardiosphere-derived cells bioactivity

CDC senescence seems more determinant than cardiac tissue donorś chronological age on CDCś proliferative and migrative properties in culture. An example of CDCś properties from a chronologically aged donor (70-years old) but with “low” senescence rate in culture (9.1%) vs. a chronologically young donor (7-months old) but with “high” senescence (14.8%) is shown in Figure 1a. “Low” senescent CDCs from the aged donor are highly proliferative, with a higher proportion of Ki-67+ cells, and with a higher ability for wound closure compared to the “highly” senescent CDCs from the young donor.

**Figure 1.**
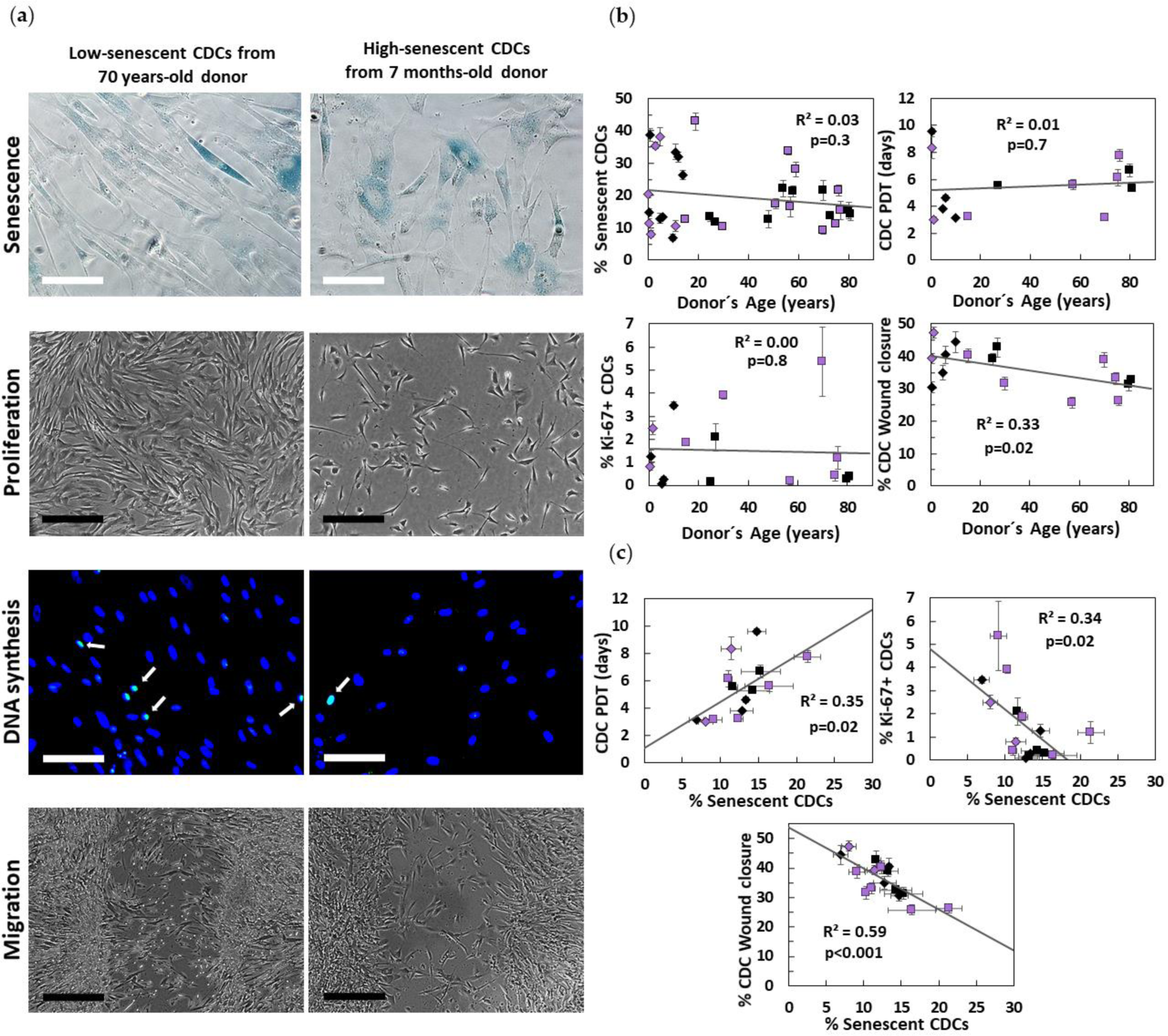
Donor chronological age vs. CDC biological age effect on different CDC properties. (**a**) Representative images from one chronologically aged donor (70 years old), but biologically young (9.1% of senescent CDCs) vs. one chronologically young donor (7 months old), but biologically aged (14.8% of senescent CDCs) after 72 hours of culture. (**b**) Chronological age is not significantly related to CDC senescence, population doubling time (PDT) or DNA synthesis capacity (% Ki-67+ CDCs) after 72 hours of culture. Chronological age significantly affects CDC migration capacity after 24 hours of wound generation. (**c**) CDC senescence however relates to PDT, DNA synthesis and migration capacity significantly and to a higher extent. Adult samples are represented with a square, while pediatric samples are represented with a diamond. Male samples are shown in black and female samples in purple. White scale bar corresponds to 100 µm, black scale bar to 400 µm.

The chronological age or sex of the 34 human donors of cardiac tissue in our study (see methods section) have shown to be unrelated to molecular markers of cellular ageing in explant-derived cells (EDCs: precursor of CDCs) and CDCs derived from tissue samples under the same conditions. The chronological age does not correlate to the genetic expression of *CDKN1A*, *CDK2A* and *TP53* in EDCs, nor any of the other genes explored (R^2^ < 0.15, *p* > 0.1). No correlation was either observed between chronological age and EDCś or CDCś senescence (Figure 1b). Donor’s chronological age did not influence CDC proliferative capacity (PDT), DNA synthesis nor VEGF secretion (R^2^ < 0.15, *p* > 0.1), but it negatively affected CDC migration (R^2^ = 0.33, *p* < 0.05, Figure 1b). Opposite to chronological age, EDCś (data not shown) and CDCs senescence correlated to the genetic expression of some aging-associated genes (i.e. *CDK2A* with R^2^ = 0.22, p < 0.05) and to most CDCs properties: CDC proliferation (PDT, R^2^ = 0.35, p < 0.05), DNA synthesis (R^2^ = 0.34, p < 0.05) and migration (R^2^ = 0.59, p < 0.001) as shown in Figure 1c. CDC senescence did not correlate to CDC telomere length or VEGF secretion (R^2^ < 0.15, *p* > 0.1).

### Scoring the anti-ageing potency of CDC-EVs in vitro

CDC-EVs were purified from CDCs obtained from 18 human donors and tested on CSCs from two human donors (target 1 with “moderate” basal senescence and target 2 with “high” basal senescence) and on endothelial cells (ECs). CDC-EVs from different donors reduced senescence of target 1 between −3.2 and −8.1 %. Similarly, in target 2, CSC senescence was reduced by all CDC-EVs as well (between −2.0 and −14.5 %). However, the anti-senescent effect of the CDC-EVs did not correlate to any of the CDC characterization parameters explored (chronological age of the donors, cardiosphere size, percentage of senescent EDCs or percentage of senescent CDCs, Figure 2). Regarding angiogenic potential, all CDC-EVs except one, significantly improved endothelial branching length (between +15 and +47 %). CDC-EVs pro-angiogenic effect did not correlate to their anti-senescent effect nor to any of the CDC characterization parameters explored (R^2^ < 0.15, *p* > 0.1).

**Figure 2.**
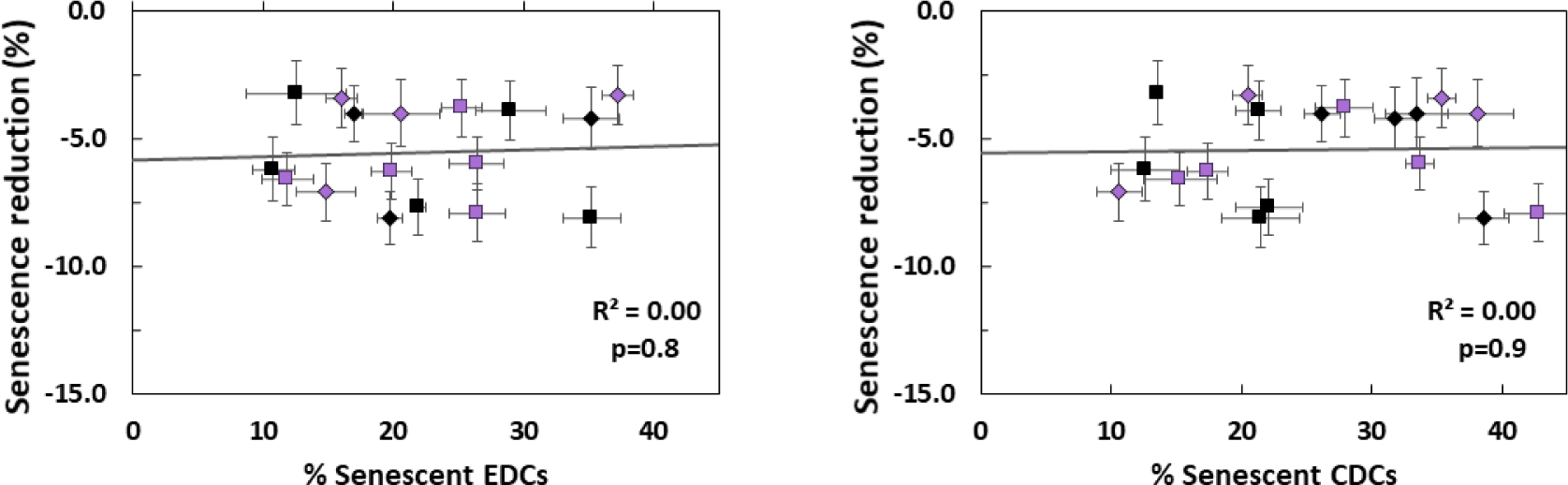
Correlation between EDC senescence and CDC senescence vs. derived CDC-EVs anti-senescent potency in one of the CSC target donors (*n* = 1199 ± 27 cells/condition). Adult samples are represented with a square, while pediatric samples are represented with a diamond. Male samples are shown in black and female samples in purple.

All CDC-EVs significantly increased CSC IL-6 secretion and most CDC-EVs reduced CSC *CDKN1A* and *CDKN2A* expression but only a few CDC-EVs reduced *TGFB1* and *TP53* expression in CSCs (Figure 3a).

**Figure 3.**
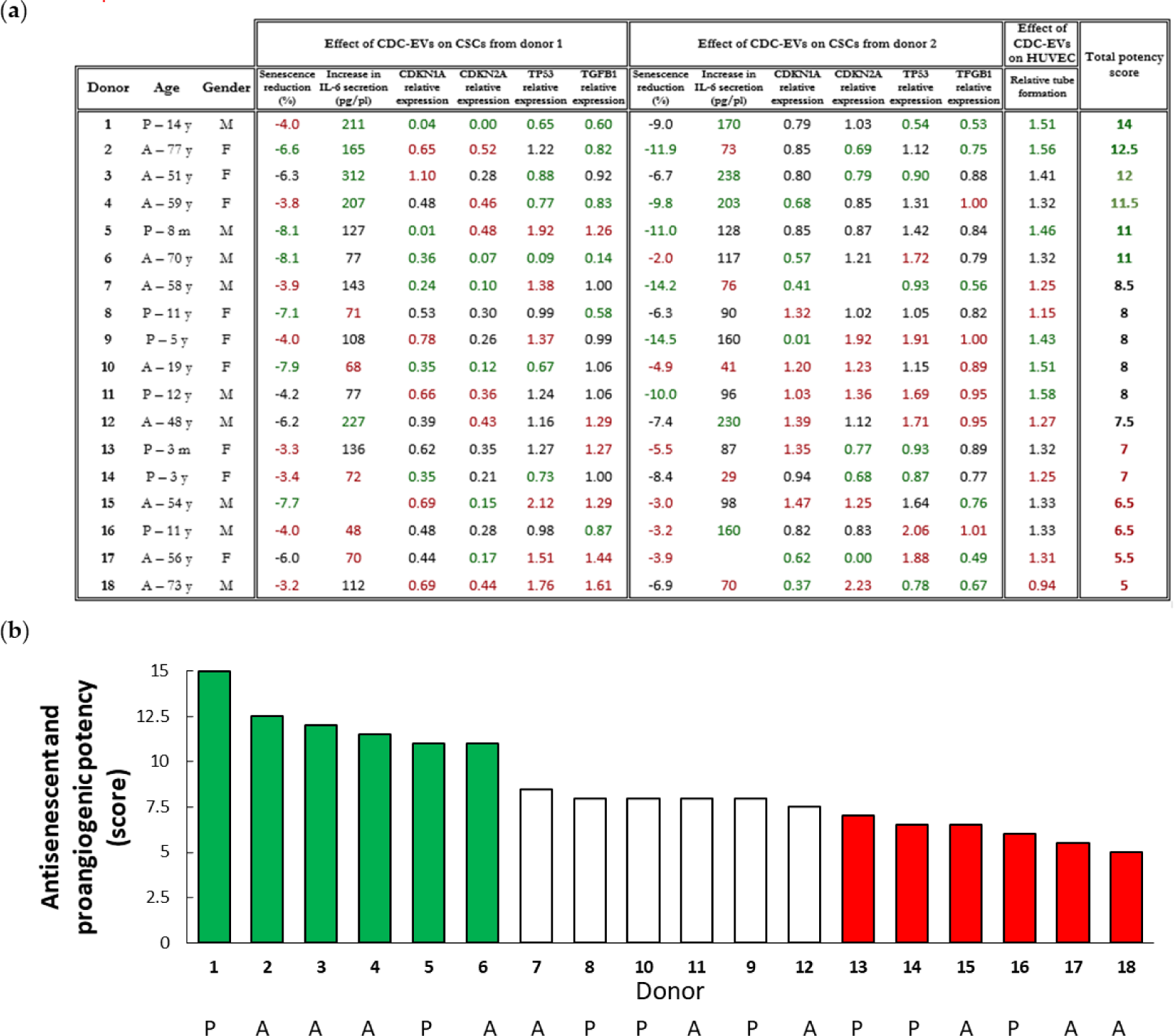
CDC-EV potency assay results for the different donors (**a**) Characterization and performance of CDC-EVs from the different donors in the different tests in the potency matrix-assay. Donor position in the potency test, age, gender, performance in the different tests in the potency assay matrix, and total potency score obtained. Reduction in CSC senescence (in two target donors) induced by the different CDC-EVs (*n* = 1199 ± 27 cells/condition in target donor 1 and *n* = 1520 ± 18 cells/condition in target donor 2). Increment in CSC IL-6 secretion vs. serum-free media (SFM) alone (in two target donors) induced by the different CDC-EVs (*n* = 3 samples/condition). *CDKN1A, CDKN2A, TP53* and *TGFB1* relative expression in CSC (from two target donors) under the effect of the different CDC-EVs (*n* = 3 samples/condition). Relative tube formation in endothelial cells (HUVEC) under the effect of the different CDC-EVs (*n* = 19 ± 1 images/condition). All data are with respect to untreated CSCs or endothelial cells under SFM. (**b**) Total score obtained in the potency matrix-assay, used for the classification of potent CDC-EVs, mild-potent CDC-EVs and non-potent CDC-EVs. A: Adult. P: Pediatric. M: Male. F: Female.

Based on the individual results obtained for the 7 ageing-related parameters evaluated in the assay, the total potency score was calculated for each CDC-EVs from different donors which were then classified as more-potent (green, total score between 11 and 14), mild-potent (white, total score between 7.5 and 8.5) and less-potent (red, between 5 and 7) (Figure 3b). The CDC-EVs with the highest potency (1 and 2, with 14 and 12.5 points, respectively, in the potency score over max. 18 possible), were among the 33 % CDC-EVs with the “best” results in most of the tests evaluated in the assay matrix. On the other hand, the least potent donor (18, with 5 points in the potency score), was among the “worst” 33 % CDC-EVs in most of the potency tests in the assay matrix, not having shown a significant beneficial effect in most of the *in vitro* tests. The performance of each of the different CDC-EVs on the different parameters of the potency matrix assay is shown in Figure 3a.

According to these results, CDC-EVs from donors 1 (14-years old, male) and 2 (77-years old, female) were selected as the more-potent and used for treating animals assigned to the potent EVs group (P-EVs) *in vivo*. CDC-EVs from donor 18 (73 years old, male) were considered as the less-potent and were used for treating the animals assigned to the less-potent EVs group (NP-EVs).

### Validation of the in vitro ageing-reversal potency assay in a rat model of premature cardiac senescence

More- and less-potent CDC-EVs selected according to their potency score in the matrix assay were tested in an *in vivo* model of premature cardiac ageing (Figure 4a). Administration of daily D-galactose (D-GAL) to rats had been demonstrated to induce cardiac-ageing (Bo-Htay et al., 2018). In our study, senescence of the cardiac tissue, measured by the expression of the *GLB1* gene, was significantly increased in rats with the daily administration of D-GAL during 20 weeks (1.56 ± 0.17 vs. 1.00 ± 0.1 in the control group, *p* < 0.05) as shown in Figure 4b. Treatment with P-EVs abrogated the D-GAL-induced ageing, since the relative *GLB1* expression in P-EV-rats was significantly lower vs. the D-GAL group (0.93 ± 0.19, *p* < 0.05) and similar to healthy controls. The use of NP-EVs showed only a non-significant trend for improvement.

**Figure 4.**
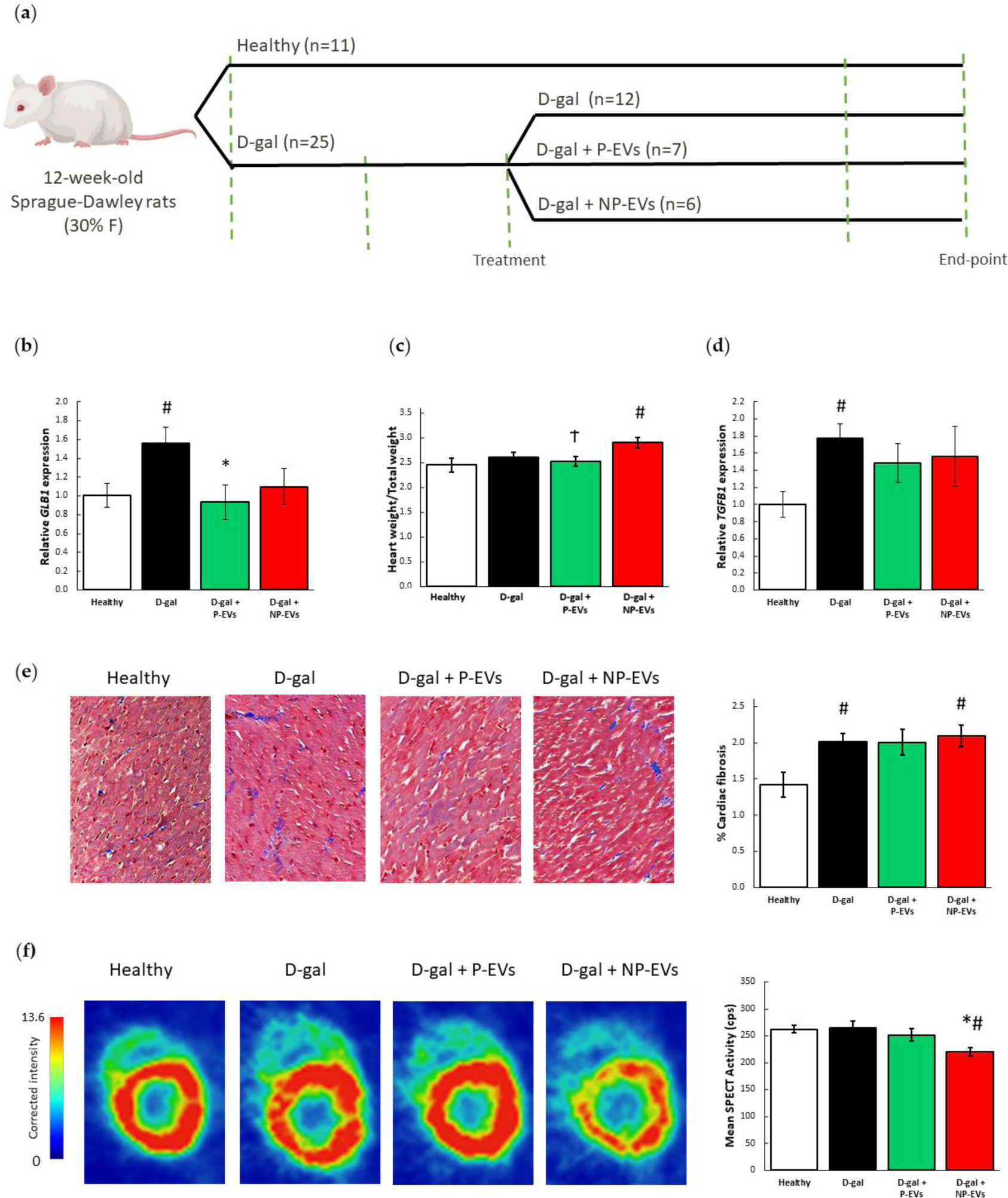
Cardiac effects of D-gal and P- and NP-EV administration. (**a**) P-EVs and NP-EVs rejuvenating and pro-angiogenic potency was tested in a rodent model of cardiac aging. Timeline including group distribution, product administration and tests performed. (**b**) Effects on cardiac senescence at genetic level (healthy: n = 6, D-gal: n = 11, D-gal + P-EVs: n = 7, D-gal + NP-EVs: n = 6). (**c**) Effects on cardiac hypertrophy on the whole population (healthy: n = 8, D-gal: n = 12, D-gal + P-EVs: n = 7, D-gal + NP-EVs: n = 6). (**d**) Effects on cardiac fibrosis at genetic (healthy: n = 6, D-gal: n = 11, D-gal + P-EVs: n = 7, D-gal + NP-EVs: n = 6) and (**e**) tissue level (healthy: n = 4, D-gal: n = 12, D-gal + P-EVs: n = 7, D-gal + NP-EVs: n = 6). Representative images of interstitial fibrosis in animals in each group. (**f**) Effects on cardiac perfusion (healthy: n = 4, D-gal: n = 4, D-gal + P-EVs: n = 6, D-gal + NP-EVs: n = 4). Representative images of Single Photon Emission Computer Tomography (SPECT) intensity corrected by injected activity, animal weight and acquisition time in animals in the different groups. F: Female. D-gal: D-galactose. EVs: Extracellular vesicles. P: potent. NP: non-potent. \**p* < 0.05 vs. D-gal, # *p* < 0.05 vs. Healthy, Ϯ *p* < 0.05 vs. NP-EVs.

**Figure 5.**
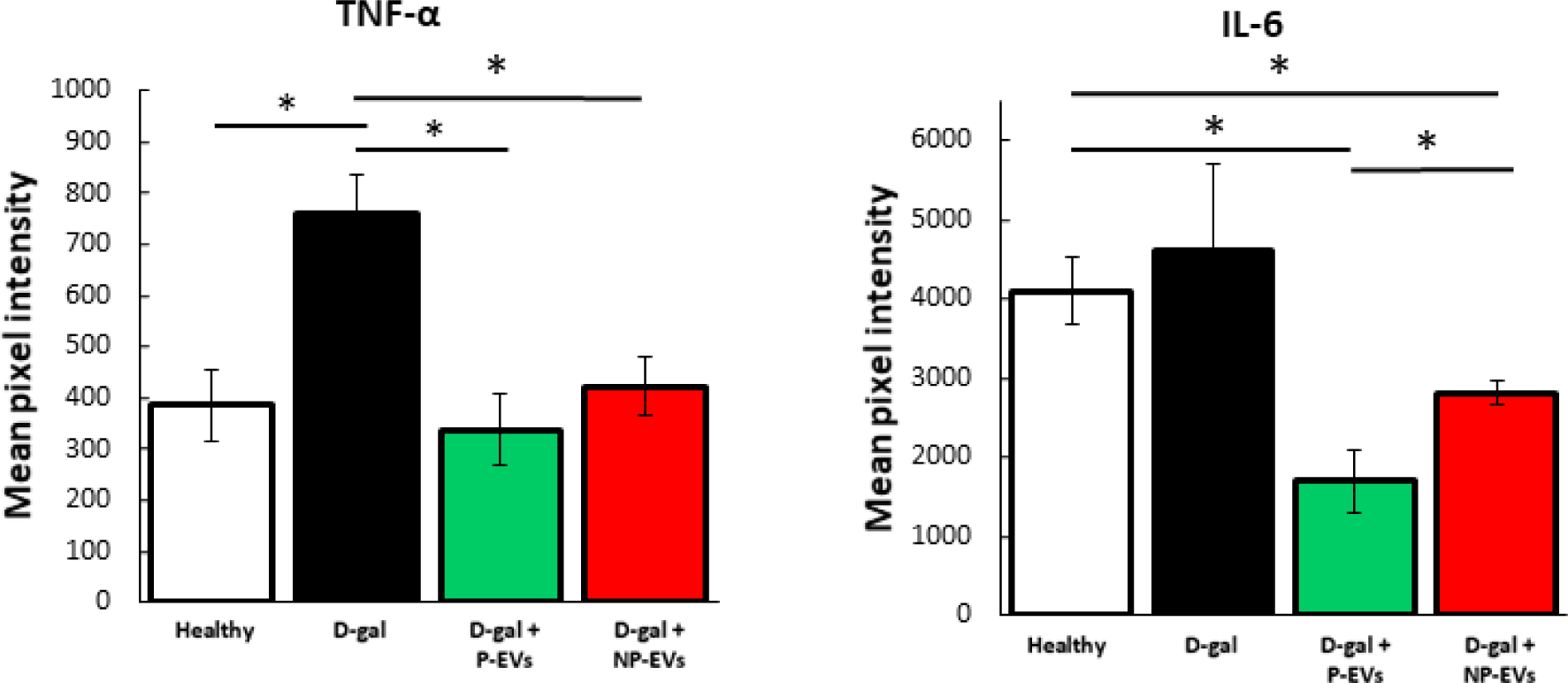
Effect of of D-gal and P- and NP-EV administration in TNF-α and IL-6 levels in peripheral blood (healthy: *n* = 4, D-gal: *n* = 4, D-gal + P-EVs: *n* = 4, D-gal + NP-EVs: *n* = 4). D-gal: D-galactose. EVs: Extracellular vesicles. P: potent. NP: non-potent. * *p* < 0.05.

Cardiac hypertrophy, measured as the ratio of the heart to the body weight, was barely increased by daily administration of D-gal in the whole group of animals (2.6 ± 0.09 vs. 2.5 ± 0.1 mg/g in the control group, Figure 4c) although the effect was more notorious in males (2.5 ± 0.05 vs. 2.2 ± 0.1 g/kg, p < 0.01; in D-GAL vs. controls respectively). Given the absence of hypertrophy with D-GAL there were no room for further improvement using P-EVs, however, rats treated with NP-EVs developed cardiac hypertrophy (2.9 ± 0.11 mg/g) with *p* < 0.05 both vs. the healthy and the P-EVs groups. When considering only males, this difference was also significant vs. the D-GAL group (*p* < 0.05).

Cardiac fibrosis was increased in D-GAL rats (Figure 4d-e). At genetic level, administration of EVs prevented from significant increase in *TGFB1* expression observed in D-GAL group (1.8 ± 0.2, *p* = 0.01 vs. healthy rats) at slightly lesser extent with NP-EVs (1.6 ± 0.3) than with P-EVs (1.5 ± 0.23) although the difference between both was not significant. Histological evaluation showed that interstitial fibrosis was significantly increased by D-GAL administration (from 1.4 ± 0.2 to 2.0 ± 0.1 %, *p* < 0.05) and in the D-GAL rats treated with NP-EVs (2.1 ± 0.1 %, p<0.05 vs. healthy). The increase of the fibrosis in rats treated with P-EVs was not significant (2.0 ± 0.2%, *p* > 0.05). Representative images of cardiac fibrosis in animals in the different groups are shown in Figure 4e.

At functional level, D-gal administration did not affect cardiac perfusion. Cardiac perfusion measured by SPECT mean activity at the endpoint was similar in the healthy, D-gal and P-EVs groups (262 ± 7, 264 ± 13 and 252 ± 12 cps respectively, all differences not significant, *p* > 0.05, Figure 4f). Nevertheless, cardiac perfusion of animals treated with NP-EVs was considerably worse than in animals in healthy and D-GAL groups (220 ± 7 cps, *p* < 0.05 vs. healthy and D-gal). Representative images of SPECT activity in animals in all groups at endpoint are shown in Figure 4f. In a randomly selected subgroup of P-EV (n=4) and NP-EV (n=2) animals SPECT was also performed at week-12, before the treatment was administered, showing an opposite trend in the perfusion evolution from week-12 to the endpoint in both groups (Suppl. Fig 3). Serum levels of inflammaging-related TNF-α and IL-6 were increased in D-GAL rats and were lower in EV-treated animal, with a slightly better profile in P-EV vs. NP-EV groups (Figure).

## Discussion

A simple *in vitro* assay proposed in this study permits to select therapeutically more potent CDC-EVs to reverse cardiac ageing in a living organism. The rejuvenating potency of CDC-EVs in our study was determined in a step-forward approach, by their ability to favorably modulate six senescence-related genetic, protein and bioactivity markers gathered in a matrix assay. Markers selected in the assay are linked to the potential MoA by which CDC-EVs are expected to exert their beneficial effects in the treatment of cardiac ageing (Valieva et al., 2022; Scheller et al., 2011; Xing et al., 1998; López-Domínguez et al., 2021; He & Sharpless, 2017; Mijit et al., 2020; Tominaga & Suzuki, 2019).

The lack of suitable *in vitro* assays to monitor the therapeutic potential of EVs currently restricts their application in clinical studies. Although challenging because of the complexity of the mode of action of native EVs, testing their biological activity by using a potency test *in vitro* or *in vivo* is recommended by expert working groups in the field (Silva et al., 2021; Grigorian-Shamagian et al., 2021). These assays will ensure that only EV-products of consistent potency are released for clinical testing or therapeutic use. In fact, as our results show some of the EVs may even have negative effects contributing to the controversial or neutral results observed in other studies (Madonna et al., 2016; Bolli et al., 2022; Smith et al., 2007; Carr et al., 2011; Grigorian-Shamagian et al., 2017; Kasai-Brunswick et al., 2017; Zhao et al., 2018). Thus, development of potency tests will promote rigorous, robust clinical testing of EV-based drug products and accelerate clinical translation. As suggested previously (Gómez-Cid et al., 2021), in this study we designed the potency test considering the anti-senescent effect of the CDC-EVs as the most relevant MoA to counteract the ageing-related cardiac changes underlying some types of heart failure with preserved ejection fraction (HFpEF). The reduction of the senescence in different cardiac cell types, as well as the proangiogenesis have been associated to multiple therapeutical benefits in cardiac ageing (Lähteenvuo & Rosenzweig, 2012; Lewis-McDougall et al., 2019; Walaszczyk et al., 2019; Yan et al., 2021).

Clinical trials in patients with HFpEF have been failing to demonstrate robust efficacy of different pharmacological strategies during decades until recently, when two drugs from the group of sodium-glucose cotransporter type 2 inhibitors proved to meaningfully reduce the HF hospitalizations and cardiovascular death (Solomon et al., 2022). However, because of the increasing prevalence of the HFpEF, with high morbidity and mortality rates, this syndrome still constitutes an important target for the new drug-development research.

In light of the increasing awareness for treatments in patients with HFpEF, the development of corresponding animal models would be highly desired (Mishra et al., 2019). There is increasing evidence that the chronic exposure to D-Galactose in rodents is associated with an enhanced cardiac expression of senescence markers, oxidative stress, inflammation, mitochondrial dysfunction, fibrosis, hypertrophy that ultimately are translated into diastolic dysfunction (Cai et al., 2022; Bo-Htay et al., 2018; Wu et al., 2017) observed in ageing-related HFpEF. By using the D-GAL model of accelerated ageing in our study we could reproduce some of the previously described effects on cardiac tissue such as an increased senescence and fibrosis and higher serum levels of inflammatory markers. However, our study failed to demonstrate an increase in cardiac hypertrophy in the global group (it was only reproduced in males), parameters of diastolic dysfunction such as end-diastolic pressure, min dP/dT, or Tau (data not shown). In terms of cardiac fibrosis, it is worthy to note that even increased in D-GAL rats compared to healthy controls, it was meaningfully lower (2 % vs. ∼8 % or higher) than the reported in other studies where CDCs and CDC-EVs showed rejuvenating effects (Grigorian Shamagian et al., 2023). All these considerations are important to take into account while interpreting at some extent only modest beneficial effects of the P-EVs given the limited room for improvement they had in our D-GAL animals.

Several limitations of our study need to be considered. As mentioned before, the rat model of D-GAL-induced premature ageing we used in the study did not recapitulate all the cardiac changes, especially those related with diastolic dysfunction, described previously (Chang et al., 2016; Hubesch et al., 2022; Kim et al., 2023; Werner et al., 2018). This might have affected mostly the potentially beneficial effects of the P-EVs. However, we did confirm our hypothesis by demonstrating a more favorable therapeutic profile of the P-EVs vs. NP-EVs in other genetic and structural parameters where the model worked appropriately, although the degree of these changes was still far from the described in naturally-aged animals (Chen et al., 2018; Grigorian-Shamagian et al., 2017; Liao et al., 2015; Liu et al., 2021). On the other hand, although the number of human donors included in our study was not small (n=18) and the chronological age range was wide (3 months - 81 years old), larger sample size, and inclusion of neonate samples, would permit to make a more robust conclusion about the implication of the chronological age of the donors on the potency of the therapeutic product, absent in this study. A larger sample size would also help defining pass and fail quantitative ranges for the therapeutic EVs instead of qualitative ones used in this study. Finally, all tissue samples were obtained from sick donors who underwent cardiac surgery for different reasons, the inclusion of healthy donors would be of interest for the purpose of the study.

In spite of all the limitations, the results highlight the importance of performing adequate potency tests before using biological products in the clinical scenario, as the use of non-potent or suboptimal products may lead to unforeseen results.

In conclusion, if confirmed in other ageing-related disease animal models, the matrix assay proposed here could be a suitable *in vitro* potency test for discerning suitable EVs for their allogenic use in clinical trials of ageing related HF-pEF.

## Acknowledgement

This research was funded by the Instituto de Salud Carlos III, Ministerio de Ciencia e Innovación, Spain: PI16/01123; PI19/00161; PI22.00792; Red de Terapia Celular, Tercel, (RD16.0011.0029) and CIBERCV (CB16.11.00292).

## Authors’ contributions

L. Gómez-Cid processed the biopsies, performed the experiments, analyzed and interpreted the data, prepared the figures and drafted the manuscript. M Cervera-Negueruela, A. Campo-Fonseca, A. Ocampo and S. Suárez-Sancho provided technical assistance. A. G. Pinto and JM. Gil-Jaurena contributed to sample collection. F Fernández-Avilés and J Bermejo acquired the funding, supervised the project and critically revised the work. Lilian Grigorian-Shamagian designed, supervised, validated the work and helped drafting the manuscript. All authors read and approved the final manuscript.

## Supplementary Figures

**Suppl. Figure 1.**
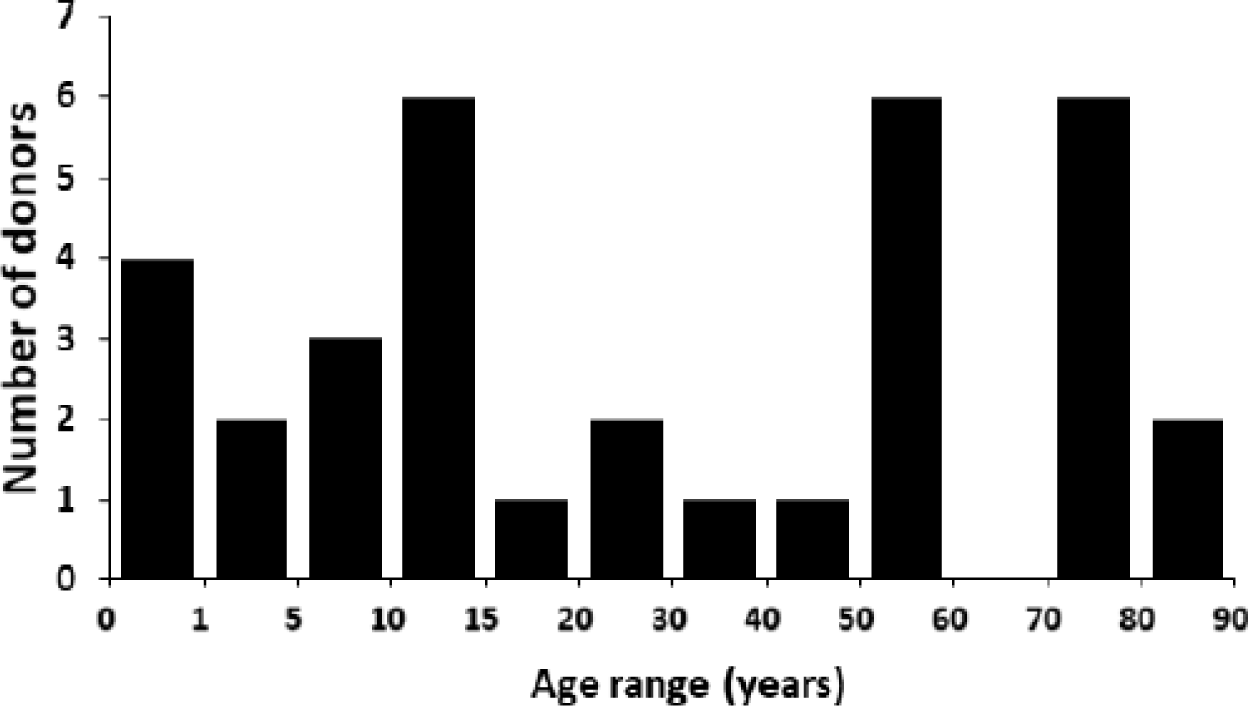
Cardiac biopsies donor distribution by age.

**Suppl. Figure 2.**
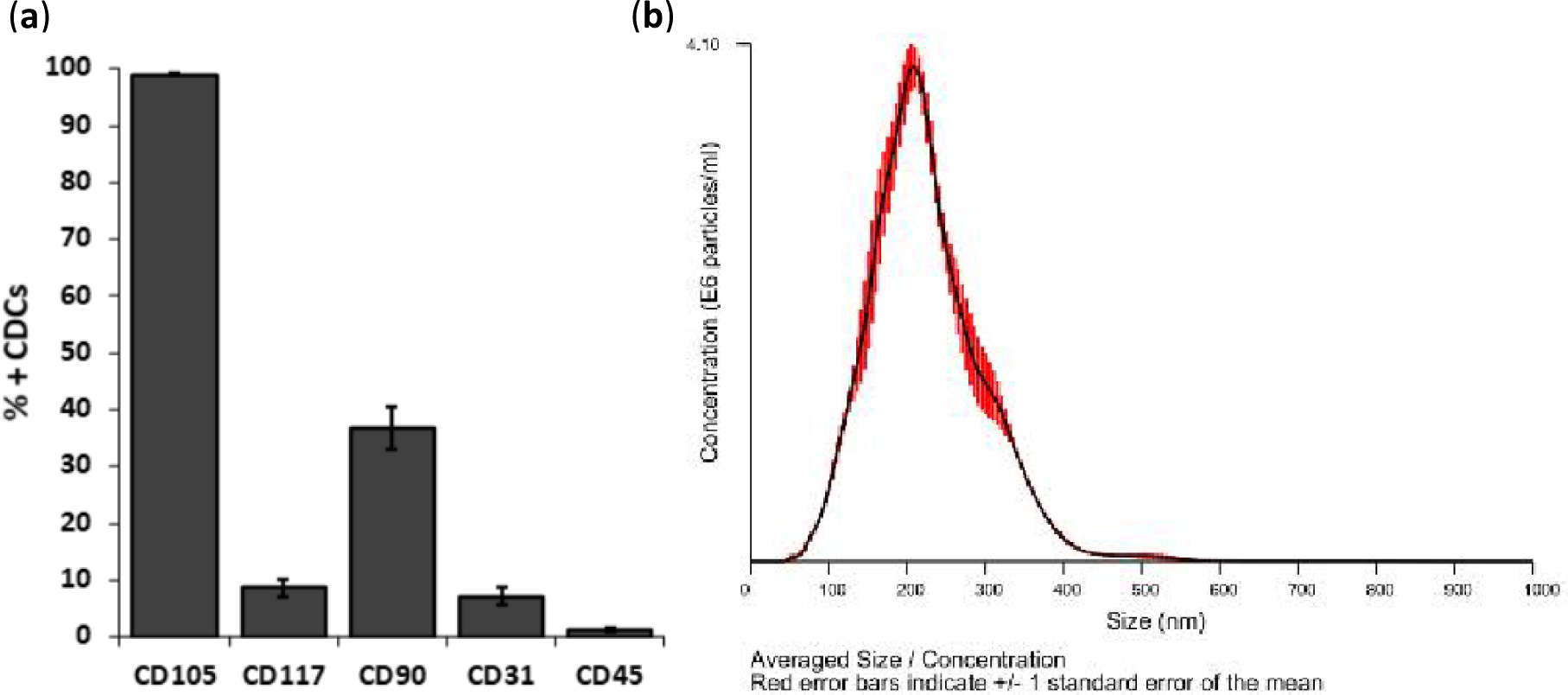
CDC and CDC-EV characterization. (**a**) Percentage of CDCs expressing CD105, CD117, CD90, CD31 and CD45 by flow cytometry (n = 18 donors). (**b**) CDC-EV quantification was performed through nanoparticle tracking analysis (NTA). Average particle size was 200 nm and maximum particle size 450 nm.

**Suppl. Figure 3.**
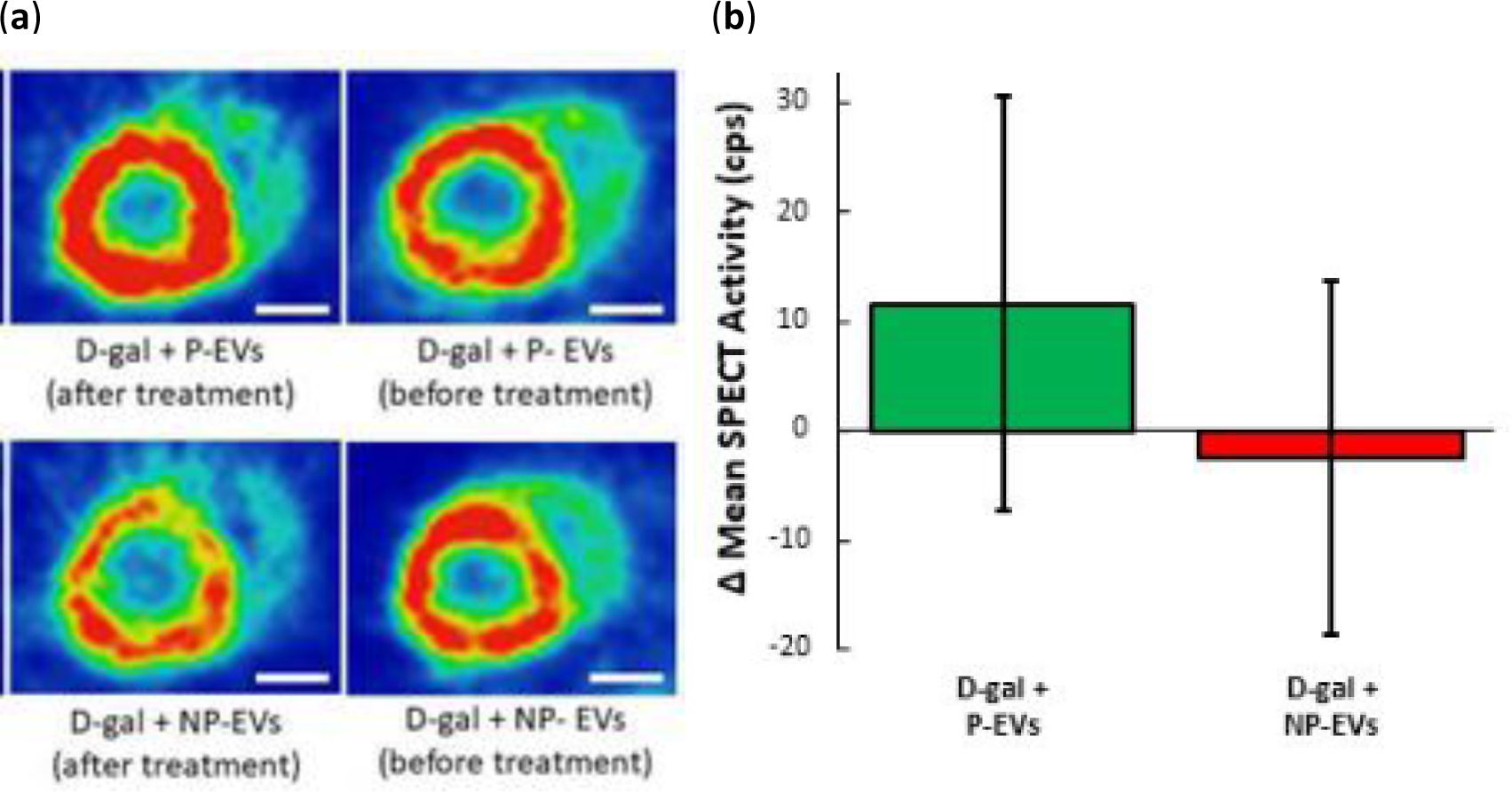
Evolution in cardiac perfusion (D-gal + P-EVs: *n* = 4, D-gal + NP-EVs: *n* = 2). (**a**) Representative images of Single Photon Emission Computer Tomography (SPECT) and intensity corrected by injected activity, animal weight and acquisition time in animals in the P- and NP-EVs groups before receiving the EVs to illustrate the effect of the treatment administration. (**b**) Evolution in Mean SPECT Activity in the two different groups. D-gal: D-galactose. EVs: Extracellular vesicles. P: potent. NP: non-potent. White scale bar corresponds to 5 mm.

## Supplementary Tables

**Suppl. Table 1.** Cardiac biopsies donor distribution by age group and sex.

|  | Males | Females | Total |
| --- | --- | --- | --- |
| *PEDIATRIC (< 14 years)* | 8 | 6 | 14 |
| *ADULTS* | 9 | 11 | 20 |
| *TOTAL* | 17 | 17 | 34 |

**Suppl. Table 2.**
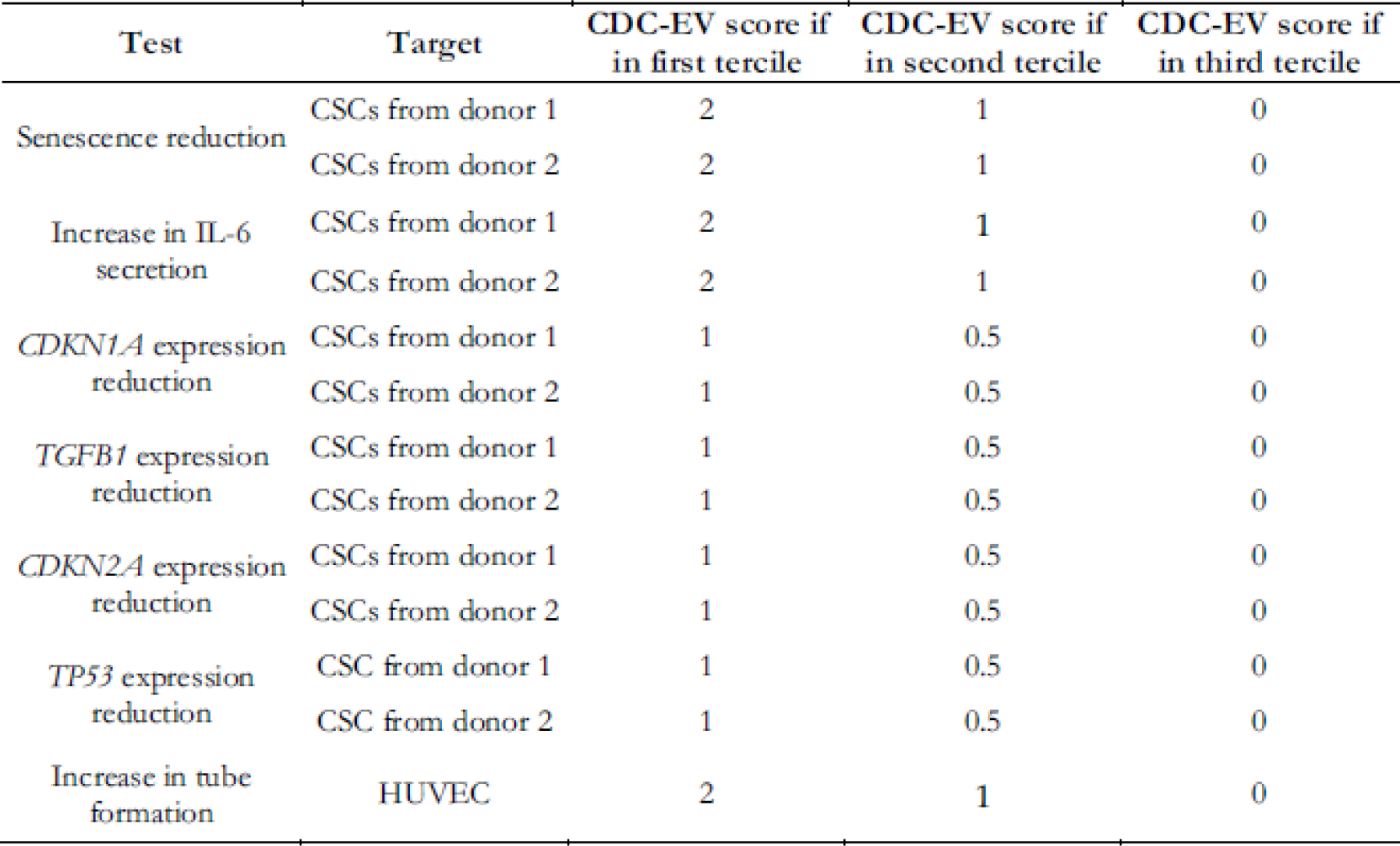
CDC-EV score assigned according to the extent of the beneficial effect in each of the different tests in the matrix-assay. The final potency score was calculated as the sum of all of them.

